# Inferring community assembly processes from functional seed trait variation along temperature gradient

**DOI:** 10.1101/2021.12.17.473108

**Authors:** Rosbakh Sergey, Chalmandrier Loïc, Phartyal Shyam, Poschlod Peter

**Affiliations:** Ecology and Conservation Biology, University of Regensburg, D-93040 Regensburg, Germany; Theoretical Ecology, Faculty of Biology and Pre-Clinical Medicine, University of Regensburg, D-93040 Regensburg, Germany; Centre for Integrative Ecology, University of Canterbury, School of Biological Sciences, Christchurch Canterbury 8041, New Zealand; Nalanda University, School of Ecology and Environment Studies, Rajgir, India

**Author notes:** Corresponding author: Sergey Rosbakh.

**Keywords:** assembly, convergence, community, competition, dispersal, divergence, facilitation, filtering, germination, seed, trait

## Abstract

1. Assembly of plant communities has long been scrutinized through the lens of trait-based ecology. Studies generally analyze functional traits related to the vegetative growth, survival and resource acquisition and thus ignore how ecological processes may affect plants at other stages of their lifecycle, particularly when seeds disperse, persist in soil and germinate.
2. Here, we analyzed an extensive data set of 16 traits for 167 species measured in-situ in 36 grasslands located along an elevational gradient and compared the impact of abiotic filtering, biotic interactions and dispersal on traits reflecting different trait categories: plant vegetative growth, germination, dispersal, and seed morphology. For each community, we quantified community weighted mean (CWM) and functional diversity (FD) for all traits and established their relationships to mean annual temperature.
3. The seed traits were weakly correlated to vegetative traits and thus constituted independent axes of plant phenotypical variation that were affected differently by the ecological processes considered. Abiotic filtering impacted mostly the vegetative traits and to a lesser extent on seed germination and morphological traits. Increasing low-temperature stress towards colder sites selected for short-stature, slow-growing and frost-tolerant species that produce small quantity of smaller seeds with higher degree of dormancy, high temperature requirements for germination and comparatively low germination speed.
4. Biotic interactions, specifically competition in the lowlands and facilitation in uplands, also filtered certain functional traits in the study communities. The benign climate in lowlands promoted plant with competitive strategies including fast growth and resource acquisition (vegetative growth traits) and early and fast germination (germination traits), whereas the effects of facilitation on the vegetative and germination traits were cancelled out by the strong abiotic filtering.
5. The changes in the main dispersal vector from zoochory to anemochory along the gradient strongly affected the dispersal and the seed morphological trait structure of the communities. Specifically, stronger vertical turbulence and moderate warm-upwinds combined with low grazing intensity selected for light and non-round shaped seeds with lower terminal velocity and endozoochorous potential.
6. *Synthesis*. We clearly demonstrate that, in addition to vegetation traits, seed traits can substantially contribute to functional structuring of plant communities along environmental gradients. Thus, the, hard’ seed traits related to germination and dispersal are critical to detect multiple, complex community assembly rules. Consequently, such traits should be included in core lists of plant traits and, when applicable, be incorporated into analysis of community assembly.

## Introduction

Now more than ever, ecologists are striving to understand the processes shaping the structure and composition of biological communities along ecological gradients (Cody et al. 1975; Cornwell und Ackerly 2009; Weiher und Keddy 1999). Disentangling the community assembly rules, i.e. the ecological processes selecting for or against species from the regional species pool and determining local community composition (Keddy 1992), is crucial both for explaining current biodiversity patterns and for predicting their future changes over the course of global change (D’Amen et al. 2017; Götzenberger et al. 2012; Newbold 2018).

In this context, trait□based approaches provide fundamental tools allowing a better understanding of community assembly processes (McGill et al. 2006). These approaches assume that functional traits, that is ‘morphological, physiological, phenological or behavioral characteristics impacting individuals’ fitness via their effects on growth, reproduction and survival’ (Violle et al. 2007) mediate species’ ecological niches and interactions (Pillar et al. 2021). Therefore, community functional trait composition should reflect the outcome of key assembly rules: abiotic filtering, dispersal and biotic interactions (Bello et al. 2013; Spasojevic und Suding 2012).

The common agreement is that multiple traits related to different organs and/or ontogenetic stages (Dayrell et al. 2018; Kleyer und Minden 2015) should be considered, because they relate to different ecological niche axes (Craine et al. 2012; Grime et al. 1997; Laughlin 2014; Leishman und Westoby 1992). This is particularly important when predicting community assembly along complex environmental gradients (e.g. elevation, latitude), as traits evolve in response to various abiotic and biotic conditions and therefore might have multiple functions (Kergunteuil et al. 2018). Yet a closer look at the published literature reveals that the prevailing majority of the examined employed traits relate to vegetative growth, survival and resource acquisition (e.g. leaf, root and whole-plant traits), while regeneration and dispersal traits are rarely considered (E□Vojtkó et al. 2020; Jiménez□Alfaro et al. 2016; Rosbakh et al. 2018). Thus, including floral, gametophyte, seed and seedling traits, which provide additional, unique information about different plant functions, in the current research agenda is intended to maximize our understanding of trait-based community assembly (E□Vojtkó et al. 2020; Jiménez□Alfaro et al. 2016; Larson und Funk 2016; Laughlin 2014; Poschlod et al. 2013; Rosbakh et al. 2018).

Ecological theory predicts that a plant species cannot be a part of the community if it possesses seed traits that are not optimised to the processes ruling the local community assembly process (Grubb 1977; Poschlod et al. 2013). Even if the environmental conditions are suitable for adult plant growth and survival (‘adult niche’ *sensu* Grubb 1977), the species’ long-term persistence in the community is ultimately defined by its ability to produce viable seeds that can successfully disperse, persist, germinate and produce vital seedlings (‘regeneration niche’; Grubb 1977). Several studies, albeit very scarce, have demonstrated that variability in seed traits related to dispersal, germination and seedling establishment is not random in plant communities (e.g. (Fernández-Pascual et al. 2017; Lebrija-Trejos et al. 2010; Rosbakh et al. 2020c; Soons et al. 2017). This suggests that seed traits are, similarly to adult plant traits, important for explaining co-existence patterns and assembly rules in plant communities (Poschlod et al. 2013). However, this remains largely unverified empirically. Surprisingly, community seed trait composition has been assessed only using a relatively small number of traits mostly related to either dispersal (e.g. (Johnson et al. 2018) or establishment (e.g. (Rosbakh et al. 2020c), thus neglecting germination and seed persistence in soil (Poschlod et al. 1998). Moreover, these are mainly ‘soft’ traits, easy-to-measure plant characteristics with low predictive power (Weiher und Keddy 1999), which carry several, often independent, functions. For example, seed mass, one of the most frequently studied seed traits in functional ecology (Larson und Funk 2016), can reflect abiotic filtering effects on germination (large seedlings from large seeds perform better at the germination stage; (Leishman und Westoby 1994), but also persistence in the soil (smaller seeds persist longer in soil; (Hodkinson et al. 1998)).

In this paper, we explore the role of vegetative growth, germination (establishment), dispersal and seed morphological traits in the assembly of mountain grasslands. We analyzed an extensive data set of 16 traits for 167 species measured in 36 grasslands located along a steep temperature gradient. Specifically, we tested whether variation in two key metrics of community functional structure (community-weighted trait mean values (CWM) and functional diversities (FD); (Ricotta und Moretti 2011)) was significantly driven by the steep temperature gradient.

First, based on previous research in similar settings (Bello et al. 2013; Chalmandrier et al. 2017), the increase in low-temperature stress as site altitude rose was expected to cause a significant shift in plant strategies towards stress-adaptation and a stronger selection of maladapted species. This strong abiotic filtering at high elevation should lead to a shift in the CWMs along the temperature gradient and a stronger trait convergence (low FD) in upland communities compared to their lowland counterparts. Because ambient temperatures control growth and development in seedlings and adult plants in a similar manner, we anticipated that abiotic filtering would have comparable effects on both vegetative and seed germination traits. Specifically, for the seed traits we expected the upland communities to be dominated by species with a risk-averse regeneration strategy aiming to minimize risks of seedling establishment in cold climates (e.g. dormant seeds with a high temperature requirement for germination and slow germination speed (Fernández-Pascual et al. 2021; Rosbakh und Poschlod 2015). Given that the high levels of low-temperature stress at high elevations (short growth period with overall low temperatures coupled with frequent and severe frost events) negatively affect seed production resulting in low seed quality and quantity (e.g. (Inouye 2008; Steinacher und Wagner 2013)), we further expected alpine grasslands to be assembled by species with comparatively lighter seeds and lower seed productivity.

Second, we expected to observe patterns congruent with the stress gradient hypothesis, which states that biotic interactions in plant communities change with the level of abiotic stress (Maestre et al. 2009). The comparatively benign conditions seen in low-elevation grasslands should promote hierarchical competition or niche partitioning, resulting in trait convergence or divergence, respectively (Münkemüller et al. 2020). Conversely, higher levels of facilitation in colder sites were anticipated to result in trait divergence (McIntire und Fajardo 2014). The shift from competition in lowland grasslands to facilitation in alpine ones should be manifested not only in vegetative traits, but also in traits related to seed germination, as seedling establishment can be strongly affected by biotic interactions with adult plants (e.g. (Kos und Poschlod 2007; Thompson und Grime 1983). In particular, we anticipated high functional divergence in seed germination traits because of temporal and spatial niche partitioning under high levels of competition, which in our study system are confined to lowlands. There, in grasslands with dense and tall vegetation, seedlings can establish themselves in gaps (spatial niches) with comparatively low competition for available resources, especially light (Thompson und Grime 1983). Additionally, early germination, before the onset of vegetative growth of the dominant species, could also considerably increase chances for seedling establishment (Forbis 2010; Kardol et al. 2013). Alternatively, amelioration of low-temperature stress by neighbors (i.e. facilitative interactions) at high elevations (Choler et al. 2001) might lead to a large number of spatial niches available for seedling establishment resulting in a higher divergence of the seed germination traits. Seed size and production were expected to be affected by the abovementioned biotic processes in the same manner.

Third, we expected to observe shifts in seed traits congruent with the change in the main dispersal vector along the temperature gradient. The warm, lowland communities should be dominated by species better adapted to zoochory, due to the higher frequency and intensity of grazing (Pellissier et al. 2010). At the colder end of the gradient we anticipated species adapted to anemochory to dominate the communities, due to frequency and intensity of vertical wind turbulences (Tackenberg und Stöcklin 2008). Dispersal processes were further expected to affect the diversity of plant dispersal traits. We predicted that the decreasing connectivity among montane grassland as elevation increases (Körner 2007) would lead to stronger convergence in dispersal traits in upland communities due to strong abiotic filtering effects on species’ dispersal ability (i.e. only good dispersers could persist in patchy alpine vegetation). In contrast, seed dispersal traits in lowland communities were expected to be divergent, due to better habitat connectivity and a larger number of various dispersal vectors available. Finally, we anticipated that the changes in seed morphological traits at the community level would also reflect the shift in the main dispersal strategy from zoochory to anemochory. Specifically, upland communities were expected to be assembled by species with light, elongated or flat seeds with comparatively large projection, a set of traits favoring dispersal by wind along longer distances (Fenner et al. 2005; Thomson et al. 2011).

## Materials and methods

### Study system

Field work was carried out in the eastern part of the Bavarian Alps (Germany), from 2009 to 2020. The region has the typical alpine relief, with steep mountain peaks composed of Triassic lime and dolomite rocks. The climate is typically montane with high mean annual precipitation rates (1500-2000 mm/year; (Marke et al. 2013)) and an altitudinal decrease in mean annual air temperatures with a lapse rate of ca. -0.6°C/100 m of elevation. The non-forest vegetation is largely represented by species-rich calcareous grasslands on nutrient-poor soils. In lowlands, grasslands are dominated by graminoids (e.g. *Arrhenatherum elatius, Helictotrichon pubescens, Carex flacca*) and tall forbs (e.g. *Buphthalmum salicifolium, Centaurea jacea*). As elevation increases, they are replaced by sedges (e.g. *C. firma, C. sempervirens*), dwarf shrubs (e.g. *Vaccinium vitis-idaea* and *Silene acaulis*) and short-stature herbs (e.g. *Ligusticum mutellina, Ranunculus montanus, Soldanella alpina*). Until the 1950s, the lowland grasslands were intensively grazed by domestic cattle and/or used for hay-making. Nowadays, the grasslands at elevations 600-1600 above sea level (a.s.l.) are used for, or managed by, low-intensity cattle grazing, whereas the subalpine and alpine grasslands above ca. 1700 m a.s.l. are occasionally grazed by sheep or wild ungulates such as the alpine ibex (*Capra ibex*) and chamois (*Rupicapra rupicapra*).

### Data collection

#### Plant composition and environmental characteristics of the study sites

Plant community composition was collected in 2009 in 36 grasslands located along an elevation gradient from 656 to 2363 m a.s.l. In each grassland, the vegetation was surveyed in ten 2 × 2 plots per site during the peak of the growing season, which was elevation specific. In each plot, the abundance of all vascular species was estimated based on six percentage classes: <0.1%, 1%–5%, 5%–25%, 25%–50%, 50%–75% and 75%–100%. The relative abundance of a species at a site was then calculated as the mean value of its abundance in all plots. In total, we recorded 379 species in all the sites.

#### Site temperature characteristics

Due to the remoteness and low accessibility of the study area, the only available data on site temperature conditions was mean annual air temperature at 2 meters above soil surface (MAT). The air temperature data were obtained from 20 weather stations located in the study area at elevations ranging from 360 to 1919 m a.s.l. MAT at each station was calculated for the period 2000–2008; from these data, lapse rates between elevation and MAT in the study region were calculated (−0.33 °C/100 m of elevation) to define the MAT in each studied grassland. Although MAT is criticised for its inability to reflect the complex thermal patterns of vegetation, especially above the tree-line (Scherrer et al. 2011), we found it to be a good predictor for local temperature conditions, and further weakly correlated with other ecological gradients (e.g. soil moisture and nutrients) in the study system (Rosbakh und Poschlod 2021). The MAT of the studied grasslands ranged between 1.8 and 7.3 °C.

#### Trait data

We sampled the most representative species in the local communities. More specifically, we did not consider species present in fewer than three sites and with a maximal abundance of <3% across all sites. In total, we collected vegetative and seed traits for 167 species. Species with measured trait data accounted for more than 80% of the total abundance of each community, allowing for a reliable estimate of functional diversity (Bello et al. 2013; Pakeman und Quested 2007).

Each species was sampled at the site where it was the most abundant following the approach described in Rosbakh and Poschlod (2015). Fully ripened seeds and fruits (hereafter ‘seeds’) were subsequently collected at maturity in these ‘optimal’ sites during the growing seasons 2009-2020. Because of low seed quality and/or quantity, the seeds of several species were collected from populations located close to the ‘optimal’ sites. Seeds were collected from a large number of randomly chosen individuals growing at a distance from each other. After collection, seeds were air-dried for several days, cleaned of flower/fruit debris, and kept dry in a cold room at 4 °C prior to the trait measurements. If not specified otherwise, the trait measurements followed the standardized protocols (Kleyer et al. 2008; Perez-Harguindeguy et al. 2016). The trait values within species were considered to be ‘fixed’, i.e. a single mean trait value was assigned per species measured under the species’ optimal ecological conditions. Therefore, our study does not consider potential intraspecific trait variation.

#### Vegetative traits

To compare the detected community assembly rules for the seed traits to those for vegetative traits, we also measured canopy height (CH), specific leaf area (SLA), leaf nitrogen content (LeafN) and foliar frost sensitivity (FoliarFS) for each species. All measurements were made *in situ* in the ‘optimal’ growing sites as described above.

#### Seed germination traits

We considered dormancy type, initial temperature of germination, germination speed and germination synchrony as the traits that represent different aspects of the seed regeneration strategy. Every species was characterized in terms of type of seed dormancy (*sensu* (Baskin und Baskin 2004)) based on published data in (Baskin und Baskin 2014; Rosbakh et al. 2020a; Rosbakh und Poschlod 2015) and the authors’ unpublished data on the seed germination ecology of the study species. Dormancy type is a categorical variable with six classes: non-dormancy (ND), physical dormancy (PY), physiological dormancy (PD), morphological dormancy (MD), morphophysiological dormancy (MPD, i.e. MD+PD), and combinational dormancy (CD, i.e. PY+PD). The seeds of a high proportion (82%) of the study species were categorized as dormant, of which 74% had a component of PD that was further subcategorized into three levels (non-deep, intermediate and deep) depending on the depth of dormancy (Baskin und Baskin 2014). Only 7% of study species possessed PY and none of them had MD or CD. Thus, to make the species comparable with each other, a relative weighted score was given to each species based on the type/depth of dormancy. Ultimately, this led to the species’ allocation to five categorical variables: ND, PY, non-deep PD/MPD, intermediate PD/MPD, and deep PD/MPD with a relative rank score of 0, 0.25, 0.5, 0.75 and 1, respectively (DormRank).

The remaining seed germination traits were obtained from germination experiments under controlled conditions (see (Rosbakh und Poschlod 2015) for further details). In brief: seeds with PD were either cold moist stratified at 4°C for six weeks or treated with 0.1% gibberellin acid prior to the germination experiment to alleviate the dormancy. Seeds of eight species were mechanically scarified to overcome physical dormancy. Seeds were germinated for 6 weeks along a temperature gradient (10/2, 14/6, 18/10, 22/14, 26/18, 30/22 °C, 14/10h photoperiod), which represents the germination conditions (from cool spring to warm summer season) in the plant communities along the elevation gradient that the seeds may encounter. Seed germination was scored regularly; the viability of non-germinated seeds was checked through inspection of embryos. The seed germination data were used to calculate:

1. minimal temperature of seed germination (T5): lowest temperature at which 5 % of all seeds germinated (Rosbakh und Poschlod 2015);
2. germination speed calculated as mean germination time in days (MGT; (Lozano□Isla et al. 2019)) with lower values reflecting faster seed germination;
3. germination synchrony calculated as the germination synchronization index (Lozano□Isla et al. 2019), with values close to one indicating that germination of all seeds occurs at the same time (more synchronized germination), while values close to zero indicate that seed germination of at least two seeds occurred at a different time (less synchronized germination).

#### Seed dispersal traits

We assumed that anemochory, epi- and endo-zoochory are the most important seed dispersal vectors in the study system (Pellissier et al. 2010; Poschlod et al. 2005; Tackenberg und Stöcklin 2008). We used seed terminal velocity (TV; m/s) as a proxy for species’ capacity for dispersal by wind, with lower values indicating longer dispersal distances. TV was calculated from hand stopped falling times from a height of 2 m with a correction for the initial acceleration phase. The height of 2 m was chosen to minimize the effect of reaction time on the hand stopped falling times (Tackenberg und Stöcklin 2008).

Epizoochory was estimated as seed attachment potential to the fur of cattle and sheep (EpiCow and EpiSheep, %), two of the most common domestic grazers in the study system. The values range from 0 to 100%, with higher values indicating potentially longer dispersal distances by the corresponding animals (Römermann et al. 2005). EpiSheep values correlate closely with seed attachment potential to ibex and chamois fur (S. Rosbakh, unpublished results) and can thus be used as proxies for epizoochory by these two species.

Species capacity for endozoochory (EndoZoo) is based on dung germinating seed surveys in the study area (Poschlod and Rosbakh, unpublished results) and published data on comparable systems (Albert et al. 2015). It is a semi-quantitative variable with three levels: 1 – seed endozoochorous dispersal is frequent (viable seeds are found in more than half of all samples) and/or in large numbers (>100 germinable seeds/kg of dry dung); 0.5 – endozoochory is rare (viable seeds are present in less than 50% of dung samples) and/or in small numbers (<100 germinable seeds per kg of dry dung); 0 – viable seeds are not found in the dung samples.

#### Seed morphological traits

Seed mass (SeedMass, mg), seed shape (SeedShape) and seed projection (SeedPr; mm^2^) were also included in our study to provide additional mechanistic explanations of seed dispersal trait variability along the MAT gradient. Seed mass (SeedMass) is the average mass of a single seed extrapolated from the weights of three samples of 100 seeds each. Seed shape (SeedShape) is the variance of seed length, width and height; it is a dimensionless value that varies between zero in perfectly round and 0.2 in disk- or needle-shaped seeds (Knevel 2005). Seed projection, a one-side are of a seed, was measured by scanning seeds on a flatbed scanner with resolution of 1200 dpi followed by seed area measurements with the help of ImageJ software (Schneider et al. 2012). Although, strictly speaking, seed production (average number of seeds produced per ramet of 10 randomly selected individuals; SeedN, pcs) is not a morphological trait, it was included in this category for the sake of brevity.

#### Data analysis

To estimate and visualize the relationship between studied plant traits, we analyzed the correlation matrix between functional traits using the package *corrplot* (Wei et al. 2017) and conducted a principal component analysis (PCA) on the species-trait matrix.

To analyze the change in the functional structure of communities, we estimated community weighted mean values (CWMs) and functional diversity (FD)(Ricotta und Moretti 2011). The CWM of a community is the average trait value weighted by species-relative abundance. It generally quantifies the trait value of the dominant species in a community and thus describes the dominating species’ adaptation strategy to given environmental conditions (Bello et al. 2021). FD reflects trait convergence or divergence (i.e. a decrease or increase in trait dissimilarity compared to random expectation) and is calculated using Rao’s quadratic entropy (Rao 1982).

Additionally, we tested whether observed functional diversity deviated from the null expectation that communities are a random sample of species from the regional pool. For each community, null functional diversity distributions were generated by permuting the columns (species) of the site-by-species abundance matrix. We then computed the standard effect sizes (SES) to evaluate the deviations of observed functional diversity values from random expectations. SES are calculated as the observed functional diversity value minus the mean of the null functional diversity values divided by the standard deviation of the null functional diversity values. A negative SES value indicates that functional diversity is convergent, i.e. lower than expected according to the null hypothesis. Conversely, a positive SES value indicates that functional diversity is divergent, i.e. higher than the null hypothesis would indicate. We further assessed the significance of functional diversity SES by identifying the proportion of random values that fell below the observed diversity value. If this rank value was below 0.05, functional diversity in that plot was considered significantly lower; if it was higher than 0.95, functional diversity in that plot was considered significantly higher.

The changes in community CWM and FD values with MAT were estimated with the help of a polynomial regression (e.g. FD ∼ MAT + MAT^2^) because a trait–environment relationship can be nonlinear (Bernard-Verdier et al. 2012; Kergunteuil et al. 2018). Where the quadratic term was not significant, this term was removed. Model assumptions were met in all cases. All statistical analyses were conducted in the R statistical environment (R Core Development Team 2021).

## Results

### Trait-trait relationships

Vegetative and seed traits were weakly correlated with each other (Appendix 1); the strongest correlations were between leaf N and seed projection (*r*=0.29, t=3.9, p<0.001); and canopy height and potential for endozoochory (*r*=0.24, t=3.2 p=0.002). Collinearity between seed germination and seed dispersal traits was also relatively low, with the strongest correlation between dormancy rank and potential for endozoochory at *r*=-0.22 (t=-2.9, p=0.004). Several morphological traits were significantly correlated with dispersal traits. The following were observed: a positive moderate correlation between terminal velocity and seed mass (*r*=0.44, t=6.2, p<0.001) and moderate negative correlations between terminal velocity and seed shape (*r*=-0.44, t=-6.3, p<0.001), between seed attachment potential to cow fur and terminal velocity (*r*=-0.23, t=-3.4, p<0.001), and between seed attachment potential to sheep fur and seed mass (*r*=-0.27, t=-4.2, p<0.001) and seed projection (*r*=-0.60, t=-9.7, p<0.001).

Importantly, we also detected a number of significant, from low to moderate, correlations within the group of traits, a pattern that supports our trait categorisation scheme. In the vegetative trait group, SLA was significantly, positively correlated with canopy height, leaf nitrogen and frost sensitivity (*r*=0.23, *r*= 0.33, *r*= 0.21, respectively; in all the cases p<0.005). Similarly, canopy height and foliar frost sensitivity displayed a positive, though weak, significant relationship (*r*=0.16, t=3.0, p=0.003).

In contrast, the traits within the germination/establishment trait groups showed stronger correlations. Dormancy rank was positively, moderately correlated with minimal temperature of seed germination and germination speed (*r*=0.34, *r*= 0.48, t=4.3 and 5.7 respectively; in both cases p<0.001) and negatively, weakly (*r*=-0.27, t=-3.7, p<0.001) with synchrony of germination, suggesting that dormant seeds tended to germinate under comparatively high temperatures, in a slow and asynchronous manner.

Among seed dispersal traits, terminal velocity and seed attachment potential to cattle fur were negatively correlated (*r*=-0.35, t=-3.4, p<0.001). The former trait also displayed a significant, positive correlation (*r*=0.40, t=5.5, p<0.001) with seed attachment potential to sheep fur.

### Variation of community weighted means along the temperature gradient

The community weighted means (CWM) of almost all traits changed significantly along the gradient of mean annual temperature (MAT), with differences as to the nature of the relationship and the explained variance (Figure 2). Vegetative traits showed the strongest variation with MAT; community-average values for canopy height, SLA and foliar frost sensitivity showed a sharp linear increase with temperature (R^2^ = 0.66, 0.80 and 0.82, respectively, p<0.001). The variation in CWM for leaf nitrogen content followed the same pattern; however, the amount of variance explained by this relationship was moderate (R^2^=0.25, p=0.002).

**Figure 2.**
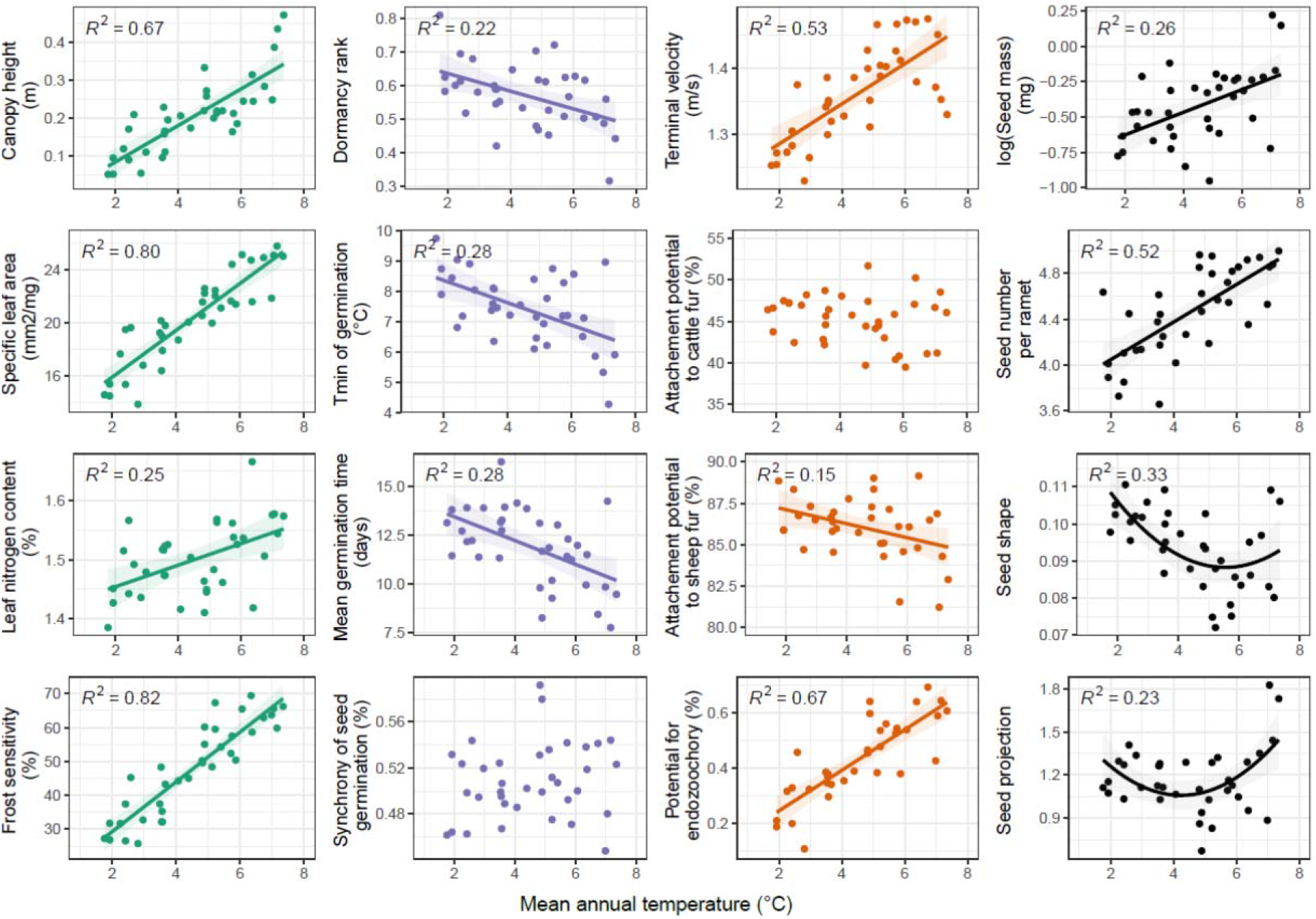
Variation in community weighted mean of 16 traits with mean annual temperatures. Full lines indicate significant relationships (p < 0.05).

As regards germination traits, the CWMs of dormancy rank, minimum germination temperature and mean germination time displayed a significant, negative relationship with the MAT gradient with a moderately similar power (R^2^=0.22, 0.28 and 0.28, respectively, all p<0.001). In ecological terms, these patterns suggest that species in upland communities tend to produce dormant seeds with a high temperature requirement for germination and relatively low germination speed. The CWM for germination synchrony did not exhibit a significant relationship with MAT gradient (Figure 1; R^2^=0.03, p=0.29).

CWM values for both seed terminal velocity and endozoochory rate increased in a sharp manner with increasing temperature (Figure 2; R^2^ = 0.62 and 0.70, respectively, both p<0.001). This pattern indicates that communities at the warm end of the gradient are dominated by species with high rates of endozoochory, while communities at the cold end of the gradient are dominated by species with high rates of anemochory. The CWM for the sheep epizoochory rate was high across the entire temperature gradient (86% on average) but significantly and weakly decreased with increasing MAT (Figure 1; R^2^ =0.15, p=0.02), suggesting a moderately lower potential for sheep epizoochory in lowland plant communities. The CWM for cow epizoochory did not vary significantly along the temperature gradient (Figure 1; R^2^=0.03, p=0.27).

The CWM for seed mass and seed number per ramet displayed a significantly negative relationship with the gradient of MAT, the strength of the former relationship being considerably stronger (Figure 2; R^2^ =0.26, p=0.003 and R^2^ =0.52, respectively, p<0.001). The CWM values for seed shape significantly but moderately increased with decreasing MAT in a non-linear manner (Figure 1; R^2^ =0.323, p=0.02). The CWM for seed projection showed a concave, significant relationship with the MAT gradient (Figure 1; R^2^=0.23, p=0.02), indicating a tendency for seeds with comparatively large surfaces to be found at both ends of the gradient.

### Variation in functional diversities along the temperature gradient

The functional diversities of vegetative traits were mostly non-random along the gradient (Figure 3). The proportion of communities with negative SES values (trait convergence) was higher for canopy height and SLA (Figure 3), whereas for LeafN and foliar frost sensitivity (FoliarFS) the majority of the communities had positive SES values (trait divergence; Figure 2). Mean standard effect size for frost sensitivity was -0.17 across communities and 16% of communities displayed significant diversity (rank below 0.05). FD of canopy height strongly decreased with decreasing MAT in a non-linear manner (Figure 3; R^2^ = 0.56, p<0.001), indicating that species in upland communities had more similar trait values than their lowland counterparts. In contrast, FD of FoliarFS showed the opposite pattern: being convergent in the lowlands, trait diversity increased towards the cold end of the MAT gradient, resulting in comparatively strong trait divergence in upland communities (Figure 3; R^2^ = 0.29, p=0.004). Furthermore, the functional diversity for SLA showed a significant, slightly concave relationship with MAT (Figure 3; R^2^ = 0.20, p=0.02), with stronger trait convergence in upland and lowland plant communities as compared to their more divergent counterparts from the middle part of the gradient. Finally, FD for Leaf N did not vary significantly across the temperature gradient (Figure 3; R^2^ = 0.12, p=0.13).

**Figure 3.**
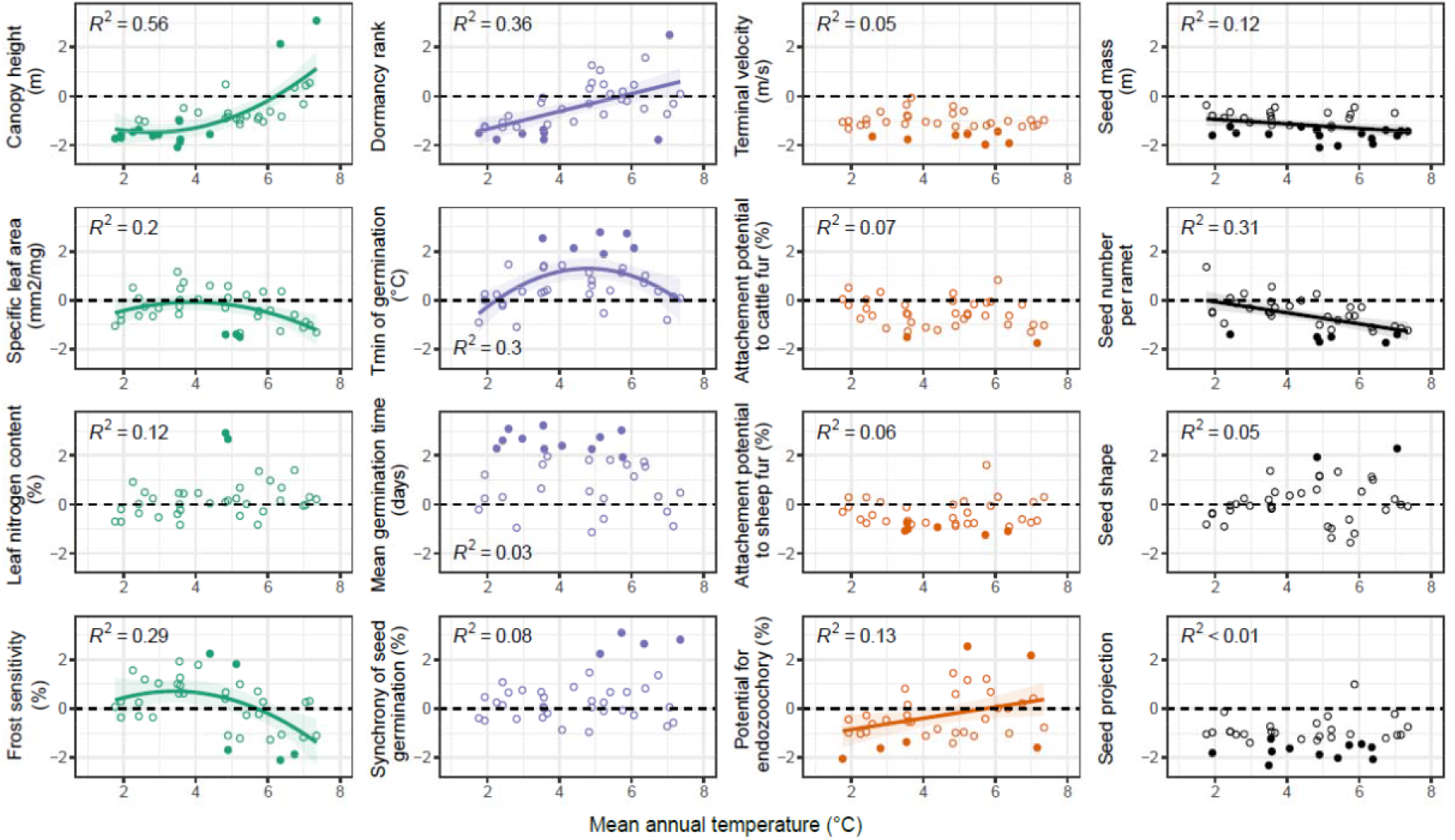
Variation in functional trait diversity (FD) of sixteen traits with mean annual temperatures. Full lines indicate that the FD-MAT relationship was significant according to the linear regressions (p < 0.05). Negative and positive standard effect size (SES) indicate a narrower and broader trait range than expected, respectively. Filled circles represent significantly low (rank lower than 0.05) or high (rank above 0.95) functional diversity.

For germination, functional diversities were mostly divergent throughout the temperature gradient (Figure 3), except for DormRank FD values that were predominantly convergent. Significant changes in FD were only found for DormRank and T5. As regards the former, SES decreased linearly with decreasing MAT (R^2^ = 0.36, p<0.001; Figure 3), changing the functional trait signature of the communities from trait divergence to convergence. As regards the latter, the FD of T5 varied moderately with MAT (R^2^ = 0.3, p=0.03; Figure 3), with more divergent communities in the middle part of the gradient as compared to its ends (i.e. concave relationship).

Dispersal trait functional diversities were almost exclusively convergent throughout the temperature gradient (Figure 3). Among dispersal traits, only the FD of EndoZoo varied significantly with MAT (negative, weak relationship; R^2^ = 0.13, p=0.03; Figure 3).

The functional diversities of seed morphological traits along the gradient of MAT followed a clear convergence pattern. FD values of SeedShape did not show any clear tendency towards convergence or divergence within the communities. MAT had a significant, positive effect on FD values of SeedMass and SeedN (R^2^ = 0.12, p=0.04, and R^2^ = 0.31, p<0.001), indicating stronger convergence of the traits in the communities in warmer sites.

## Discussion

Regeneration has always been considered a fundamental aspect of plant community ecology (Grime et al. 1981; Grubb 1977; Keddy 2010), yet it has been largely neglected in studies of community assembly (Jiménez□Alfaro et al. 2016; Poschlod et al. 2013; Saatkamp et al. 2019). In this study, we show that seed traits are only weakly linked to vegetative traits and thus constitute independent axes of plant phenotypical variation that may be affected differently by ecological processes. We thus provide the first comprehensive evidence that little-studied seed functional traits are important determinants of plant community structure in addition to vegetative functional traits. Specifically, we demonstrate that abiotic conditions, biotic interactions and dispersal acting differently on the functional traits of plants at different stages of their life cycle and ultimately shape the structure of mountain grassland communities.

### Temperature strongly filters plant communities

Confirming our expectation, abiotic conditions have a prominent impact on dominant vegetative traits of mountain grassland communities. Community weighted means of vegetative traits shifted strongly (or moderately in the case of leaf nitrogen content) along the temperature gradient (Figure 2). Specifically, the increasing low-temperature stress towards higher sites seemed to select for short-stature species (Bello et al. 2013; Bucher und Rosbakh 2021; Pellissier et al. 2010), and to further filter out tall species (based on the convergent functional diversity of height in those sites). This shows the critical importance of small stature for plants to cope with abiotic stress, as this allows heat accumulation near the ground during the short growing season (Körner 2021). The shift in the mean trait values of SLA and leaf N also mirrors the shift of leafeconomics strategies along the temperature gradient from fast-growing species with fast nutrient acquisition in warmer sites (high SLA and leaf N values) to slow-growing and nutrient-conservative species (low SLA and leaf N values in cooler sites (Rosbakh et al. 2015; Spasojevic und Suding 2012). Finally, plants were more frost resistant in colder sites, showing the importance of that trait for coping with frost events, which at higher elevations can occur even during the summer months. In contrast to plant height, the functional diversity of these traits was not convergent in cold sites, possibly indicating that despite the strong climatic stress, other ecological processes may promote the coexistence of species with different trait values (see below).

Community weighted means for three out of four seed germination traits (degree of dormancy, temperature requirement for germination and germination speed) presented a clear response to the temperature gradient, but those relationships were less strong than for vegetative growth traits (Figure 2). We attribute these shifts in CWM to the effects of abiotic filtering on germination strategies in alpine environments. Low-temperature stress (e.g. frequent, severe frost events) at the colder end of the gradient selects for plants that produce seeds with a higher degree of dormancy, high temperature requirements for germination and comparatively low germination speed (Baskin und Baskin 2014; Fernández-Pascual et al. 2021; Rosbakh und Poschlod 2015). Furthermore, there was an increasing convergence in seed dormancy with decreasing temperatures, which implies that plant communities in the colder sites are largely assembled from species with similarly high degrees of seed dormancy that are able to postpone seed germination to more favorable conditions for seedling establishment in late spring – early summer via mechanisms of seed physiological dormancy (Baskin und Baskin 2014). These adaptations aim to match seed germination timing with the most suitable environmental conditions to maximize establishment success. Specifically, longer cold stratification periods necessary to break physiological dormancy prevent seeds from germinating shortly after dispersal, whereas high base temperatures of germination and slow germination speeds postpone germination of non-dormant seeds to late spring-early summer, characterized by a comparatively low probability of severe frost events (Rosbakh et al. 2020b; Rosbakh und Poschlod 2015).

The decrease in CWMs for seed size and seed production along the lower temperature gradient suggested that species in alpine communities produce a small quantity of lighter seeds. This can be explained by the presence of harsh climatic conditions that are stressful for sexual reproduction (Figure 2). A short growth period coupled with frequent and severe frost events considerably limits pollination, fertilization and seed maturation (Lundemo und Totland 2007; Rosbakh und Poschlod 2016; Steinacher und Wagner 2013), thereby filtering out species that require more resources to produce a large quantity of heavy seeds. Alternatively, the dominance of light seeds in alpine communities could reflect the increasing role of anemochory along the elevational gradient (see below).

### Shifts in biotic interactions from lowlands to uplands filter vegetative and seed traits

The stress gradient hypothesis predicts that the increasing stress along the elevational gradient should lead to a change in biotic interactions from competition to facilitation (Maestre et al. 2009). Our results are partially congruent with this hypothesis and further suggest that in addition to abiotic filtering, biotic interactions can also filter certain functional traits in mountain plant communities.

As expected, the relatively benign climate in low-elevation grasslands promotes plants with competitive strategies including fast growth and resource acquisition (comparatively high CWM values for canopy height, SLA and leaf N; Figure 2). Likely because of the moderate trade-off between those traits and frost resistance, species from the lowlands also invest less in protecting their leaves against negative temperatures (high CWMs for frost sensitivity in lowlands; Figure 2). The convergent FD values in lowland grasslands further suggest that frost-resistant plants are excluded from these communities, most likely due to their comparatively low competitiveness (Bucher und Rosbakh 2021). In contrast, the high FD for canopy height may indicate that both small and tall plants are able to co-occur, suggesting that some degree of niche partitioning associated with this trait maintains species coexistence in those benign but competitive environments (Münkemüller et al. 2020). Surprisingly, in upland communities, the functional diversities of SLA and frost sensitivity are higher than in the lowland communities. This suggests that some species with high foliar frost sensitivity and high SLA can persist despite the strong abiotic stress. We attribute this pattern to a comparatively higher number of available thermic niches in topographically more complex higher elevations (Scherrer und Körner 2011) or to increasing facilitation among co-occurring species, with frost-resistant species creating local environmental conditions for the persistence of frost-tender species (Cavieres und Badano 2009; Choler et al. 2001).

Similarly to vegetative traits, seedling establishment in the lowlands seems to be constrained mainly by higher levels of competition. Dominant species in the lowlands tend to possess different regeneration strategies including germination after a comparatively short stratification period (low T5 community weighted mean values) and high germination speed (low MGT community weighted mean values) that allows germination in early spring before the closure of dense vegetation cover (Figure 2). Although the germination traits of dominant species tend to shift across the temperature gradient, there is strong divergent functional diversity of T5, MGT and SYN in low to mid-elevation grasslands (Figure 3). This suggests that niche partitioning stabilizes the coexistence of multiple regeneration and temporal niches (Grubb 1977, Bernard-Verdier 2012). In the uplands, this pattern vanishes and may indicate that the increasing abiotic filtering (which promotes trait convergence, see above) overrides the signal of biotic interactions.

Another interesting finding of our study is the strong convergence of FD values for seed mass and seed production throughout the entire gradient (Figure 3), a pattern that can be attributed to both abiotic and biotic filtering effects on the functional community composition structure. While in the uplands trait convergence could be related to the impact of abiotic filtering (see above), in the lowlands, the convergence of the traits might reflect the hierarchical competition favoring plants producing comparatively large seeds in large numbers, two traits necessary to ensure regeneration in dense, tall grassland vegetation (Jakobsson und Eriksson 2000; Leishman und Westoby 1994).

### Seed dispersal traits and community assembly

The relationship between community functional trait structure and the dispersal ecology of mountain vegetation is traditionally less studied than the impact of abiotic and biotic filtering. There is, however, evidence that dispersal ability determines species capacity to colonize patches of suitable habitat and maintain sink populations beyond the limits of their abiotic niche (Dullinger et al. 2012; Isabelle et al. 2012). Our results demonstrate that seed dispersal traits are also involved in community assembly. The most striking result is the strong decrease in community mean seed terminal velocity and potential for endozoochory along the temperature gradient (Figure 2). These distinct changes in dominant trait values point to a clear shift in the dispersal strategy from endozoochory in lowland sites to anemochory in higher elevation grasslands. This is likely due to the prevalence of opposite dispersal vectors at both ends of the temperature gradient. In warmer sites, the higher rate of endozoochory may be due to the long history of grazing by domestic animals (Gilck und Poschlod 2021; Poschlod 2014) that may have favored the immigration and persistence of endozoochorous plant. In colder sites, the lower terminal velocity and higher values of seed shape indicate that many plants are likely dispersed by the intense vertical turbulence (Tackenberg und Stöcklin 2008) and moderate warm upwind (Tackenberg et al. 2003a) in the upland habitats that allow effective dispersal over long distances in patchy alpine vegetation (Tackenberg et al. 2003b).

Rates of epizoochorous potential are remarkably higher for sheep than for cattle, suggesting that the former are an effective seed dispersal vector available in all grassland communities along the entire gradient (Figure 2). We also detected a weak yet significant increase in seed attachment potential to sheep fur along the temperature gradient (Figure 2). This pattern can be explained by traditional sheep grazing in upland grasslands or similar properties of sheep and wild ungulates’ fur as regards collecting and retaining seeds during dispersal (S. Rosbakh, unpublished data). Because it is mainly upland grasslands that are grazed by wild ungulates, the corresponding communities might have been assembled by species better adapted to this type of seed dispersal.

Both epizoochory rates and seed terminal velocity are correlated with seed mass, seed shape and seed projection across species. In consequence, the shift in seed dispersal strategies along the temperature gradient can also be seen in the community patterns of seed morphological traits (Figure 2). As previously shown, wind-dispersed seeds tend to be light and non-round in shape (e.g. elongated or winged seeds; Appendix 1), two traits that positively affect terminal velocity values (Fenner et al. 2005; Thomson et al. 2011). As for epizoochory by sheep, light diaspores with an elongated shape facilitate ‘anchoring’ in animal fur (Römermann et al. 2005). As seen previously, seed mass may be filtered in the lowlands by competitive interactions. The joint impact on seed mass of the competitive filter in the lowlands and the wind and epizoochory dispersal filter in the highlands may indicate why the functional diversity of seed mass, seed shape, sheep and cattle epizoochory, and terminal velocity are consistently convergent along the temperature gradient.

### Conclusions and implications

Our study reveals that in addition to vegetation traits, seed traits can also substantially contribute to functional structuring of plant communities along environmental gradients. We clearly demonstrate that a combined study of vegetative traits and seed germination, dispersal and morphology traits in plant community ecology is critical for the detection of multiple, complex community assembly rules. Specifically, we find that in montane grassland located along a steep temperature gradient 1) abiotic filtering mostly impacts vegetative traits and to a lesser extent, seed germination and morphological traits, 2) biotic interactions, specifically competition, were also found to have an effect on all types of traits but in different ways and 3) the changes in the main dispersal vector affect the structure of dispersal traits and by extension, morphological traits. These findings lend empirical support to the recent argument that plant trait research should consider multiple traits representing different ecological niche axes, i.e. different organs and/or ontogenetic stages (Craine et al. 2012; Kleyer und Minden 2015; Laughlin 2014). Therefore, along with (Jiménez□Alfaro et al. 2016; Saatkamp et al. 2019) we advocate that ‘hard’ seed traits related to germination and dispersal should be included in core lists of plant traits and, when applicable, be incorporated into analyses of community assembly.

The detected community-level seed trait responses to changing temperature have important implications not only for studies on the community assembly process, but also for land use and climate change research and conservation/restoration ecology. As regards the former, our results indicate that CWM and FD of seed traits can be effective in predicting vegetation changes in upland grassland communities. For example, the fast temperature rise in mountain ecosystems (Lamprecht et al. 2018; Rumpf et al. 2019) may lead to relaxation of low-temperature stress filtering effects on the seed dormancy, initial temperature of germination and germination speed trait values of warm-adapted species, resulting in an altered composition of communities in cold sites (i.e. increase in frequency and abundance of species from lower elevations). As regards the latter, the detected patterns in dispersal trait variation can support some nature conservation and/or restoration decisions. The functional structure and composition of lowland grasslands seem to be strongly dependent on or affected by the cattle grazing dominant here. Therefore, to conserve plant biodiversity in existing grasslands or re-establish abandoned ones, we recommend maintaining or re-introducing grazing at the historic levels, as applicable.

## Acknowledgments

We thank the numerous students and research assistants who helped with the trait measurements. SP is grateful to the Alexander von Humboldt Foundation, Germany for his Humboldt Experienced Researcher Fellowship. SR and PP were partially supported by the FORKAST project (TP12 Poschlod). LC acknowledges funding from the European Union’s Horizon 2020 research and innovation programme under Marie Skłodowska-Curie agreement No 840946 (Project “CLIMB”).

## Author contributions

SR conceived the study and performed the experiments; SR and SP compiled the dataset; SR and LC analyzed the dataset. SR and LC wrote the first version of the manuscript. All the authors helped in critically revising the manuscript and gave final approval for publication.

## Data availability statement

Should the manuscript be accepted for publication, the data will be archived at Zenodo.

## Appendix

Appendix 1. Trait-trait correlation plot and matrix (Pearson’s correlation coefficient).

Appendix 2. Principal component analysis of species-trait data.

