## Supplementary figures and images for "Inferring community assembly processes from functional seed trait variation along temperature gradient"

### Appendix 1

Appendix 1. Trait-trait correlation plot and matrix (Pearson’s correlation coefficient)

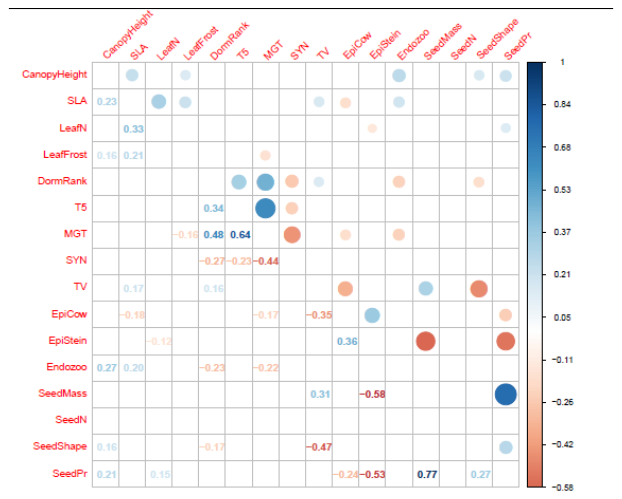
