## Appendix 2 for "Inferring community assembly processes from functional seed trait variation along temperature gradient"

The twelve seed traits studied could be summarized by two principal components (PCs) accounting together for 29.9% of the total variance. The third PC only accounted for an additional 10.8 % of total variation (Figure S1; Table S1). PC1 explained 16.2 % of the variance and loaded most heavily (Pearson’s r >|0.48|) and positively on dormancy rank, initial temperature of germination and mean germination time (Figure S1) and negatively on germination synchrony, seed attachment potentials to cow and sheep furs. Thus, PC1 reflected the main differences in seed traits of species studied and separated those characterized by dormant seeds with high temperature requirements and slow, asynchronous germination and comparatively low potential for epizoochory from species with non-dormant seeds, which can germinate over a broad range of temperature conditions in a fast and synchronous manner and have a high ability for epizoochorous dispersal.

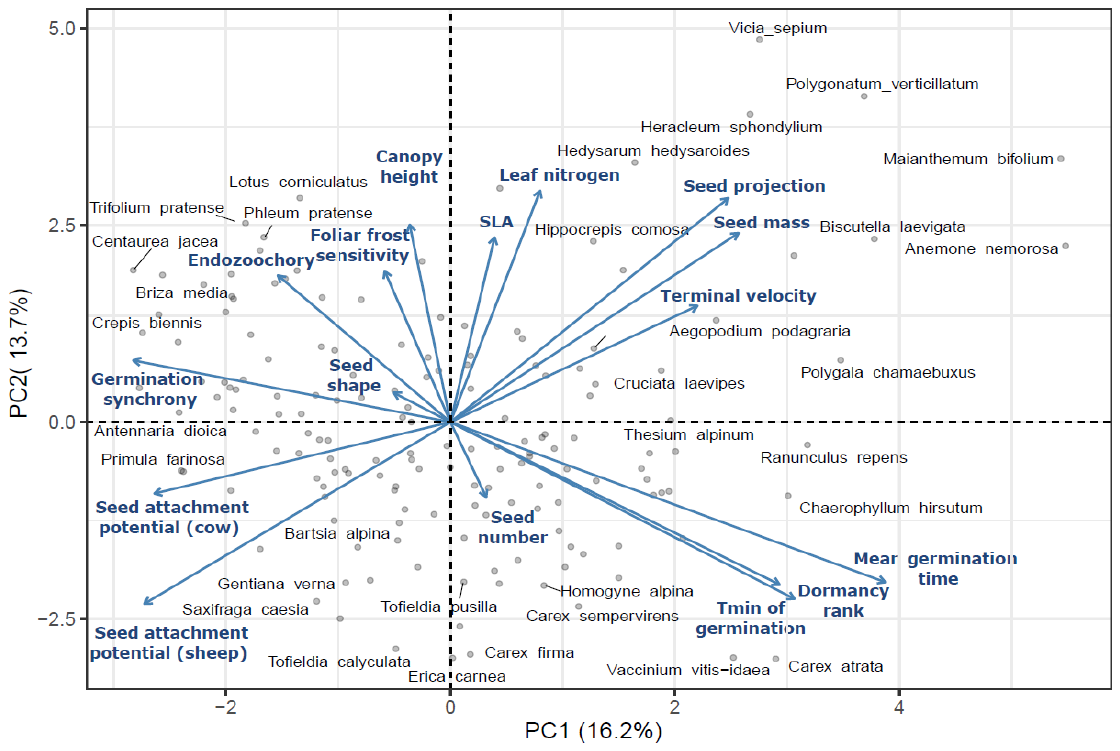

Figure S1. Two-dimensional illustration of the PCA ordination on the species-trait matrix of the studied 167 species. See Table S1 for associated eigenvalues, proportion of variance and trait loadings on first two principal components. Arrows indicate the direction of each trait loading; each point represents one of the species.

Table S1. Summary of principal components analysis (PCA) on species-trait matrix of 167 study species.

|  | **PC1** | **PC2** | **PC3** |
| --- | --- | --- | --- |
| **Eigenvalue** | 2.6 | 2.2 | 1.7 |
| **Proportion of variance** | 16.2 | 13.7 | 10.8 |
| **Cumulative proportion** | 16.2 | 29.9 | 40.7 |
| **Vegetative traits** |  |  |  |
| Canopy height | -0.07 | 0.46 | 0.31 |
| Specific leaf area | 0.07 | 0.43 | -0.15 |
| Leaf nitrogen | 0.15 | 0.54 | 0.12 |
| Foliar frost sensitivity | -0.11 | 0.35 | 0.14 |
| **Germination traits** |  |  |  |
| Dormancy rank | 0.54 | -0.38 | -0.14 |
| Tmin of germination | 0.56 | -0.41 | 0.27 |
| Mean germination time | 0.71 | -0.37 | 0.20 |
| Germination synchrony | -0.52 | 0.14 | -0.22 |
| **Dispersal traits** |  |  |  |
| Terminal velocity | 0.40 | 0.27 | -0.70 |
| Seed attachment potential (cow) | -0.48 | -0.17 | 0.10 |
| Seed attachment potential (sheep) | -0.50 | -0.42 | -0.12 |
| Endozoochory | -0.28 | 0.34 | -0.02 |
| **Seed morphological traits** |  |  |  |
| Seed mass | 0.47 | 0.44 | -0.29 |
| Seed production | 0.06 | -0.18 | -0.04 |
| Seed shape | -0.09 | 0.07 | 0.80 |
| Seed projection | 0.46 | 0.52 | 0.38 |

The second PC accounted for an additional 13.7 % of the variance and loaded most heavily (r >|0.43|)) and positively on vegetative traits including canopy height, SLA and leaf nitrogen. Additionally, seed mass (*r*=0.44) and seed projection (*r*=0.50) loaded heavily on this axis. Therefore, PC2 separated tall, fast-growing species with high foliar nitrogen contents and seed size from short-statured, slow-growing species with small leaf N values and comparatively small seeds.
